# Stoic: Fast and accurate protein stoichiometry prediction

**DOI:** 10.64898/2026.03.13.711535

**Authors:** Daniil Litvinov, Lorenzo Pantolini, Peter Škrinjar, Gerardo Tauriello, Caitlyn L. McCafferty, Benjamin D. Engel, Torsten Schwede, Janani Durairaj

## Abstract

**Motivation:** Protein complexes are central to cellular function, but experimental determination of their structures remains challenging. Structure prediction methods require prior knowledge of stoichiometry - the number of copies of each protein entity within a complex. Current approaches rely on computationally expensive brute-force methods that run structure prediction on multiple stoichiometry combinations, often with limited accuracy.

**Results:** We introduce *Stoic*, a method that uses protein language model embeddings to predict protein complex stoichiometry. Our approach learns to identify interface residues that participate in protein-protein interactions, rather than relying on global sequence features. By integrating these interface-aware embeddings into a graph neural network, *Stoic* achieves fast and accurate stoichiometry prediction for both homomeric and heteromeric targets.

**Availability:** Source code for inference and training along with web versions are available in the repository at https://github.com/PickyBinders/stoic.

## Introduction

Protein structure prediction has undergone a revolutionary transformation with approaches such as AlphaFold2 (AF2) and others (Jänes and Beltrao, 2024). However, these breakthroughs have primarily focused on individual protein chains, while many biological processes depend on protein complexes composed of multiple subunits of the same or different protein entities. Often the information about which proteins assemble into a complex is known, while stoichiometry - the number of copies of each unique protein chain is not. A critical limitation of current structure prediction methods is their requirement for this prior knowledge. Stoichiometry extends beyond individual structure prediction, since both major benchmarking efforts, CAMEO (Continuous Automated Model Evaluation) (Robin et al., 2026) and CASP16 (Critical Assessment of Structure Prediction) (Zhang et al., 2026), have established stoichiometry prediction as the first task before complex structure modelling can proceed.

Current approaches to this challenge frequently rely on running AF2-like methods on multiple stoichiometry combinations, then using confidence scores to distinguish correct quaternary states (Liu et al., 2025a; McGuffin et al., 2025; Shor and Schneidman-Duhovny, 2024). This brute-force approach is not only computationally expensive but also has limited accuracy, especially for large heteromeric targets (Elofsson, 2025). While template-based approaches like SWISS-MODEL exist (Waterhouse et al., 2018), they face significant challenges in accurately assigning stoichiometry for complexes with many individual entities, particularly where finding templates containing homologs for all entities becomes difficult.

Protein language models (pLMs) have demonstrated a remarkable ability to extract structural and functional information from sequence alone. Trained on millions of natural sequences without explicit structural labels, these models learn latent representations that capture evolutionary constraints encoding protein structure and residue conservation (Zhang et al., 2025). Recent work has shown that pLM embeddings also carry signals relevant to stoichiometry prediction: Seq2Symm (Kshirsagar et al., 2025) leverages them to predict homomeric protein symmetry, while QUEEN (Avraham et al., 2023) predicts copy numbers for homomers. However, these approaches rely on average-pooled embeddings that capture global protein properties but may miss residue-level signals essential for understanding protein-protein interactions. In addition, prediction of stoichiometry of heteromeric complexes remains unsolved and less attempted.

Here we present *Stoic*, a novel method that combines graph neural networks with interface-specific protein language model embeddings to predict protein complex stoichiometry. *Stoic* learns to identify and weight interface residues, the specific amino acids that participate in protein-protein interactions, for embedding pooling rather than relying on global sequence features. By integrating these interface-aware embeddings into a graph neural network that introduces complex-related context, *Stoic* achieves fast and accurate stoichiometry prediction not only for homo-but also heteromeric targets, while *Stoic* interface residue prediction enables interpretability.

## Methods

### Datasets

The training and benchmarking datasets used consist of biological assemblies from the Protein Data Bank (PDB) (Burley et al., 2017) after applying various filtering criteria, described in Table 1.

**Table 1.** Dataset Statistics. Number of protein complexes at different stages of filtering.

| Filtering Step | N# Complexes |
| --- | --- |
| Original PDB | 247,947 |
| Remove structures without resolution | 233,136 |
| Remove capsids | 213,817 |
| Remove proteins with helical symmetry | 213,145 |
| Remove non-protein structures | 210,433 |
| Sequence length filtering (15-1740 residues) | 199,991 |
| Deduplication | 117,956 |

For each complex, we extracted the copy number of each unique entity from its first biological unit. The combination of copy numbers for a complex (stoichiometry) served as our target variable. The “Deduplication” steps consists of sorting the data by resolution and removing duplicates based on the combination of sequences and corresponding stoichiometry, retaining the structure with the highest resolution for each unique combination. From this filtered set we extracted statistics of copy number distributions, and arrived at 14 classes (1, 2, 3, 4, 5, 6, 7, 8, 9, 10, 12, 14, 24, and 16, in the order of frequency in the dataset) representing 99.5% of the entire PDB.

The training set consists of 99,098 filtered complexes with a release date before 1 June 2023. For training the weighted pooling head, we also extract interface residues from each entity as any residue with any atom within 6Å from another entity in the complex.

To make the benchmark set, we used MMseqs2 (Steinegger and Söding, 2017) clustering with a minimum of 30% sequence identity and 80% coverage to assign a cluster label to each protein entity within each complex released after 1 June 2023, and then define the complex cluster label as the combination of the entity cluster labels. The complexes with the highest resolution from each complex cluster-stoichiometry combination were retained and further filtered to those where at least one entity has *<*30% sequence identity to any complex in the training set. Thus, the benchmark dataset consists of 1,350 diverse complexes released after 1 June 2023 and with low homology to those in the training set. This set is further divided into homomeric (one unique entity) and heteromeric (more than one unique entity) complexes.

As Seq2Symm predicts homomeric symmetry rather than copy number, we mapped prediction classes between Seq2Symm and *Stoic* (Supplementary Table S1) and constructed a subset of 396 homomers from the benchmark dataset where predictions could be compared between the two methods.

### Architecture and losses

Our model architecture integrates sequence embeddings from protein language models (pLMs) with graph neural networks to predict protein complex stoichiometry. As shown in Figure 1, *Stoic* uses ESM2-650M (Rives et al., 2021) (4-bit quantized for memory efficiency) to obtain residue-level embeddings for each unique protein entity in a complex, and then uses a learned weighted pooling mechanism to aggregate residue embeddings into a fixed-length protein embedding. The pooled embeddings act as node features in a fully connected graph, used as input to a graph convolutional neural network (GCN) which outputs protein copy numbers as node labels. We formulate this as a multi-class classification problem, with 14 copy number classes. The combination of predicted copy numbers across all input proteins represents the predicted stoichiometry.

**Figure 1.**
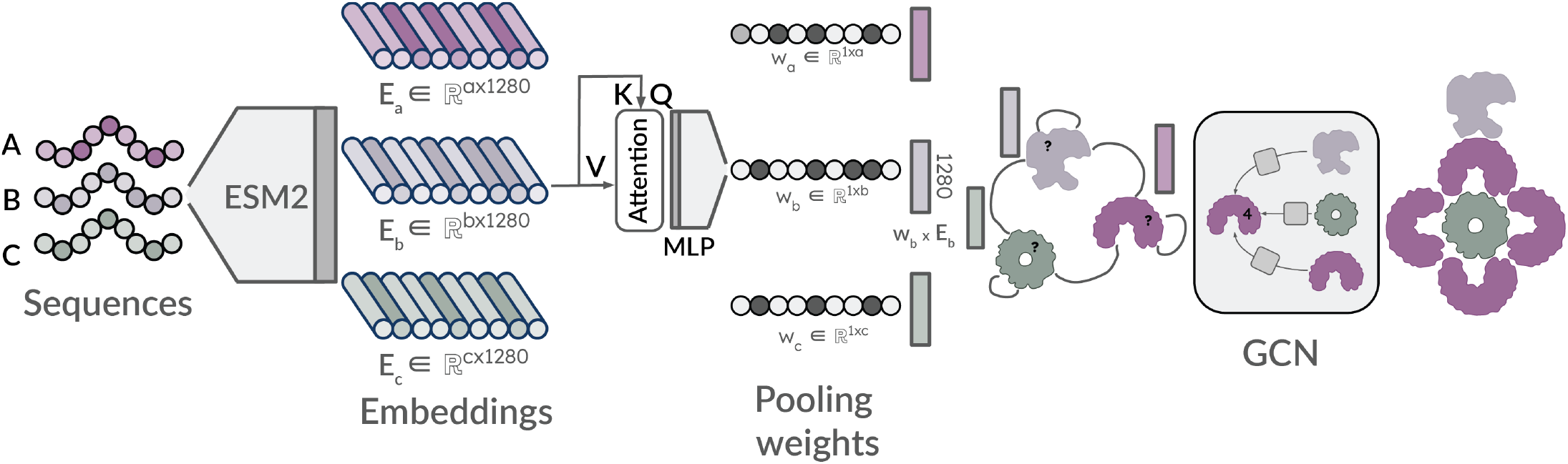
Overview of the model architecture. The architecture integrates residue-level sequence embeddings from ESM2-650M for each unique protein entity in a complex, which are aggregated into a fixed-length protein embedding using a learned weighted pooling mechanism. The pooled embeddings serve as node features in a fully connected graph, which is processed by a GCN introducing the complex context that outputs protein copy number classes as node labels. The combination of predicted copy numbers across all input proteins represents the predicted stoichiometry.

The model is trained using two complementary losses. The first (Equation 1) is the main complex loss for the copy number classification task, which steers the model to predict global stoichiometry correctly. This loss uses the sum of logarithms of individual entity cross-entropy losses between *l*_*i*_ (logits of individual entity) and *c*_*i*_ (copy number class) across all *n* entities in a complex. This is mathematically equivalent to multiplication but provides better gradient properties during training. This formulation penalizes the model more heavily for incorrectly predicting any single component of a complex compared to standard cross-entropy loss, where such errors would be obscured, especially for complexes with many entities. To counteract the effects of extreme class imbalance (e.g. class 7 and 9 make up 0.3% of the training set) we introduced class weights calculated using effective number of samples defined as 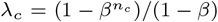, where *n*_*c*_ is the number of samples of class *c* and *β* = 0.9999 is a hyperparameter (Cui et al., 2019).

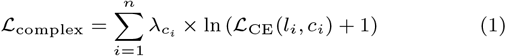

The second (Equation 2) is an auxiliary focal loss for predicting interface residues (*r*), which helps the model learn appropriate weights (*w*) for the pooling mechanism by identifying which residues are most relevant for stoichiometry prediction.

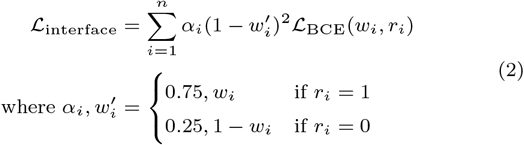

The *α* weights help account for the imbalance in the task, where most residues are not in the interface.

The overall stoichiometry for a complex is predicted as follows: for each entity, the model outputs logits for the copy number classes. These logits are converted to ordinal ranks by sorting in descending order, such that the class with the highest logit receives rank 1. Top-N stoichiometry candidates are then identified via beam search (beam width 10) over all entities in the complex, where candidate combinations are scored by the sum of their per-entity ranks. The lowest-scoring i.e., most confident combinations are returned as the top predictions.

#### Ablations

To investigate the impact of our architectural choices we train a series of models each ablating different aspects of *Stoic*. The *Naive* model simply predicts a copy number of one for all input entities. The *No-Context* model uses a multi-layer perceptron (MLP) on average-pooled ESM embeddings to predict copy number independently for each entity - i.e. this model does not make use of the information of other proteins in the complex. The *No-Weights* model instead adds the GCN component on top of the average-pooled ESM embeddings, thus incorporating heteromeric context. The *No-Auxiliary* model adds learned weighted pooling based on self-attention block to the *No-Weights* model but still only trains on the loss described in Equation 1, and thus the residue weights learned are those which improve prediction of stoichiometry but are not explicitly guided to be interface residues. The *Stoic* model on the other hand, incorporates the addition loss described in Equation 2, thus encouraging both the prediction of interface residues and subsequent use of those residues for pooling, copy number, and thus stoichiometry prediction. Finally, the *Ideal* model works with the unrealistic scenario where interface residues are already known (extracted from the ground truth) and used directly for embedding pooling. All models requiring training were trained for 100 epochs using the same training set and optimiser parameters. These include the AdamW optimizer with a base learning rate of 5*e*^−4^ and a weight decay of 0.01. A OneCycleLR scheduler was employed with a maximum learning rate of 5*e*^−4^ for the pooling parameters and 5*e*^−3^ for the remaining parts of the model. Supplementary Figure S1 describes the detailed architecture of *Stoic*.

### AlphaFold3 structure prediction

We assess the combination of *Stoic* with AlphaFold3 (Elofsson, 2025) using data from CAMEO (Robin et al., 2026) from 14 June 2025 to 15 December 2025. For each of 4,170 protein complexes, we predicted five AlphaFold3 models using a single seed and the top-predicted stoichiometry from *Stoic*. Complexes where the number of tokens exceeded the limit for an A100-40G GPU were excluded, resulting in 4090 targets. The accuracy of predicted structures was assessed using OpenStructure v2.8.0 (Studer et al., 2025) with the `compare-structures` action; iLDDT (Abramson et al., 2024) and QS-global (Bertoni et al., 2017) scores were used to estimate the effect of incorporating stoichiometry provided by *Stoic*. For 409 targets we also obtained MultiFOLD2 (McGuffin et al., 2025) predictions from CAMEO data. We compare to the CAMEO-AF3 baseline (labelled Naive) which runs AlphaFold3 with one copy of each entity in the complex.

## Results

### Context and pooling improve stoichiometry prediction

We show the power of the *Stoic* architecture through baselines and ablations on the benchmark set in Figure 2A, with the global stoichiometry prediction accuracy on homomeric complexes (in blue triangles) and heteromeric complexes (in purple squares). Note that each complex has at least one entity with *<*30% sequence identity to the training set. Our Naive baseline simply predicts a copy number of one for all entities in the complex, the current approach used in CAMEO (Robin et al., 2026) for their AlphaFold3 (Abramson et al., 2024) baseline. This is followed by an MLP on average-pooled embeddings (*No-Context*) i.e. no graph context and no weighted pooling. Architecturally, this model matches those of QUEEN (Avraham et al., 2023) and Seq2Symm (Kshirsagar et al., 2025), both designed for homomeric stoichiometry prediction where other entities are not relevant. Adding the context of the other proteins in the complex through graph convolutions strongly improves prediction performance for heteromers with a minor improvement for homomers (*No-Weights*). This mimics the concurrently released StoPred (Liu et al., 2025b), where such context was also proven to be important for accurate heteromeric stoichiometry prediction. However, adding learned weighted pooling both without (*No-Auxiliary*) and with using an interface-specific auxiliary loss (*Stoic*) further improves performance and reaches ~30 percentage points higher accuracy compared to the baseline. We also show the idealistic scenario where interface residues are obtained from the ground truth structure and used for pooling (*Ideal*), demonstrating that interface residues contain very relevant and strong signals for predicting stoichiometry, and supporting our approach to employ auxiliary interface losses to improve pooling and thus prediction performance. The *No-Auxiliary* model performs only slightly worse than *Stoic* on the global stoichiometry prediction task, indicating that the task already informs the subset of residues relevant for pooling. However, as shown in Figure 2B, the *No-Auxiliary* model does not learn to predict interface residues like *Stoic* does. Interface prediction unlocks the door to interpretability and an orthogonal measure of confidence as described in Section 3.3.

**Figure 2.**
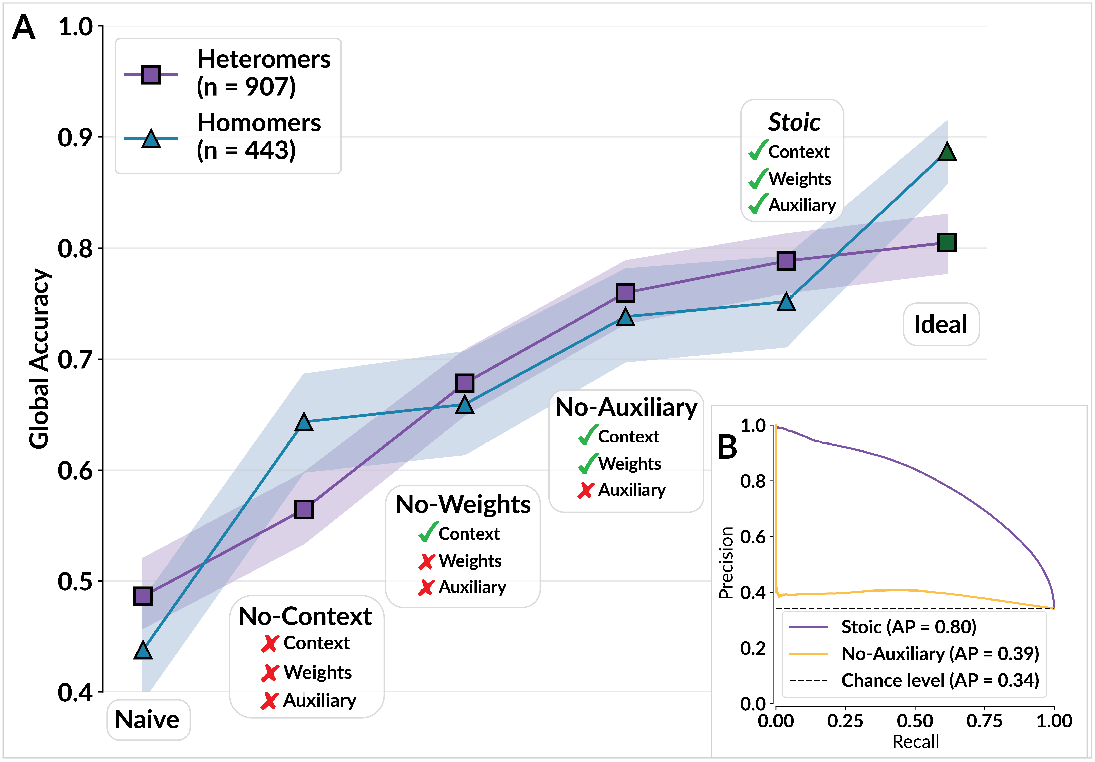
Model Ablations. **A)** Accuracy of global stoichiometry prediction for ablations removing different aspects of the network. *Naive* = predicting a copy number of 1 for all entities in a complex; *Ideal* = using the interface residues from ground truth complex structures to pool residue embeddings before the GCN step. Other ablation details are described in Section 2.2.1. **B)** Residue-level interface prediction precision-recall curve for *Stoic* and *No-Auxiliary* models, where interface residues are defined as those within 6Å of another entity.

### *Stoic* provides accurate stoichiometry predictions

The confusion matrix for node-level classification performance of copy number prediction (Figure 3A), only on proteins with *<*30% sequence identity to our training set, shows good performance across the board including for rare stoichiometries (e.g. 24 and 9). Performance remains robust across varying levels of sequence similarity to the training data, with only a moderate decrease in the lowest similarity bin (0.00–0.30), indicating that *Stoic* generalizes well to sequences distant from the training set Supplementary Figure S2. This generalisability also holds true for global stoichiometry prediction on the benchmark dataset, where a prediction is labelled correct only when all the individual protein entities within a complex have their copy numbers predicted correctly (Figure 3B). For this global task, we also compare to the template-based approach implemented in SWISS-MODEL (Waterhouse et al., 2018) which uses HHSearch (Steinegger et al., 2019) against the PDB (with a release date before 1 June 2023) followed by QSQE-based template selection (Bertoni et al., 2017). We consider a prediction as a success if any of the QSQE-selected templates has the same stoichiometry as the target (green solid bar). We also show the success rate when any of the up to 50 templates found contain the correct stoichiometry (dashed lines). For monomers and homomers where the number of unique entities is one, we compare to Seq2Symm (blue bar), which predicts symmetry for homomeric complexes (see Section 2.1 for details on mapping symmetry predictions to copy numbers). Thus, *Stoic* outperforms both template-based and previous pLM-based approaches in assigning the correct stoichiometry. The advantage over template-based approaches is especially clear in complexes with many individual entities where finding templates containing homologs for all entities becomes difficult. For both *Stoic* and Seq2Symm we also show best-of-3 predictions (dashed lines), indicating that further room for improvement in stoichiometry ranking remain and also potentially allow for cases where a protein differs in its stoichiometry based on its environment, For example, the metamorphic protein Selecase (López-Pelegrín et al., 2014) forms stoichiometries of 1, 2, or 4 depending on conditions, and *Stoic* ranks these among its top-4 predictions (4, 2, 14, and 1). On the Complex Portal dataset with experimentally determined stoichiometries, *Stoic* improves global stoichiometry accuracy over the naive baseline (Figure S3C).

**Figure 3.**
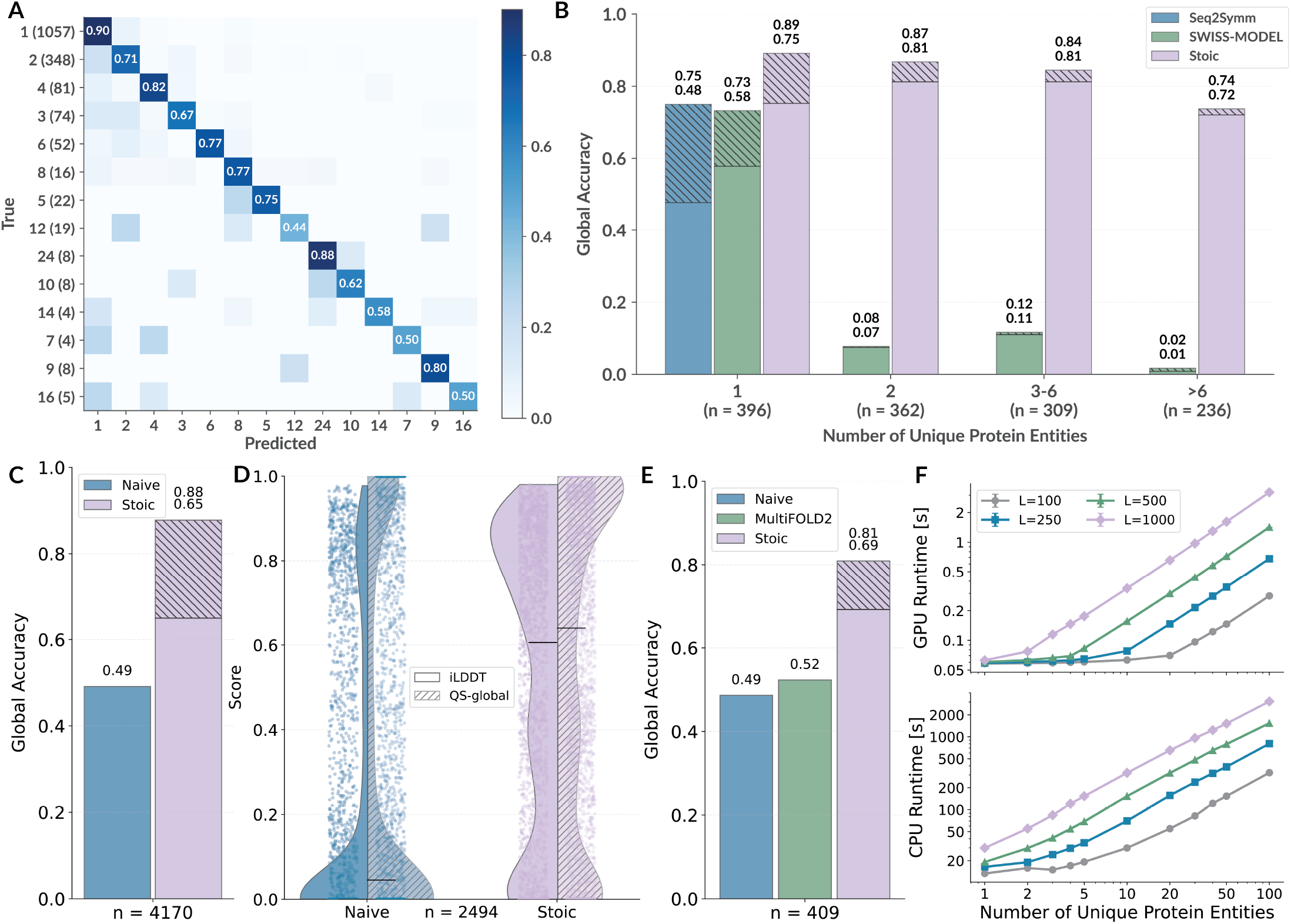
Copy number, stoichiometry and structure prediction performance. **A)** Confusion matrix of *Stoic* copy number prediction across all proteins in the benchmark set with *<*30% sequence identity to the training set. **B)** Global stoichiometry prediction accuracy across the benchmark set for Seq2Symm (blue, only for cases with one unique entity), template search (green), and *Stoic* (purple) divided across complexes with differing number of unique entities. The blue and purple dashed bars represent Seq2Symm and *Stoic* best-of-3, while for Template search the green solid bar represents SWISS-MODEL’s QSQE-based stoichiometry and the green dashed bar represents the best-of-50 templates. **C)** Global stoichiometry prediction accuracy across the CAMEO set for Naive prediction (where every entity is assigned a copy number of one) and *Stoic* (best-of-3 shown as dashed bar). **D)** Distribution of iLDDT (solid) and QS-global (dashed) scores for AlphaFold3 models generated using Naive (blue) and *Stoic* (purple) predicted stoichiometries. Note that monomers are excluded from this set as iLDDT and QS-global are not defined. **E)** Same as C but on the subset for which MultiFOLD2 (green) predictions are available in CAMEO. **F)** *Stoic* runtime measurements for different number of entities and lengths (L). All calculations were run on an A100-40G GPU and AMD EPYC 7742 64-Core CPU.

To evaluate the practical impact of accurate stoichiometry prediction on downstream structure modelling, we combined *Stoic* with AlphaFold3 (Abramson et al., 2024) on CAMEO (Robin et al., 2026) targets released between June and December 2025 (see Methods Section 2.3). The CAMEO baseline (labelled *Naive*) assigns a copy number of one to every entity, which is the default when stoichiometry is unknown. We compare this to AlphaFold3 models generated using the top-ranked stoichiometry predicted by *Stoic*. As shown in Figure 3C, *Stoic* substantially improves global stoichiometry accuracy over the Naive baseline across the CAMEO dataset. This improvement in stoichiometry directly translates to better predicted structures: Figure 3D shows that AlphaFold3 models built with *Stoic* stoichiometries achieve higher iLDDT and QS-global scores, confirming that providing the correct oligomeric state enables AlphaFold3 to produce more accurate complex models.

We further compare to MultiFOLD2 (McGuffin et al., 2025) predictions available in CAMEO for a subset of 409 targets (Figure 3E). MultiFOLD2 enumerates candidate stoichiometries by running AlphaFold2 (Jumper et al., 2021) on multiple combinations and selecting the highest-scoring result. Supplementary Figure S3A,B shows the comparison of *Stoic* + AF3 and MultiFOLD2 on the 282 non-monomeric targets for stoichiometry prediction accuracy, iLDDT and QS-global scores. We achieve better stoichiometry prediction accuracy and iLDDT and QS-global scores while requiring only a single *Stoic* forward pass followed by a single AlphaFold3 run, making it orders of magnitude faster. As shown in Figure 3F, *Stoic* inference takes under two seconds even for complexes with 100 unique entities on a single A100 GPU.

### Embeddings from interface residues inform stoichiometry

*Stoic* learns to predict residues participating in protein interfaces as a secondary task, using these intermediate predictions to generate more meaningful embeddings for the main stoichiometry prediction task. The precision-recall curve for predicting interface residues shown in Figure 2B, demonstrating good performance on this task. We showcase some examples of these residue predictions in Figure 4A, where residues with an assigned weight of *>*0.5 (the threshold yielding the best interface prediction F1-score) for one protein in each complex are highlighted. It is interesting to note that not all interface residues are relevant for stoichiometry prediction and, conversely, not all residues with high weights are in the interface. Further exploration of the residues selected by *Stoic* could be useful for interpretability studies exploring complex formation.

**Figure 4.**
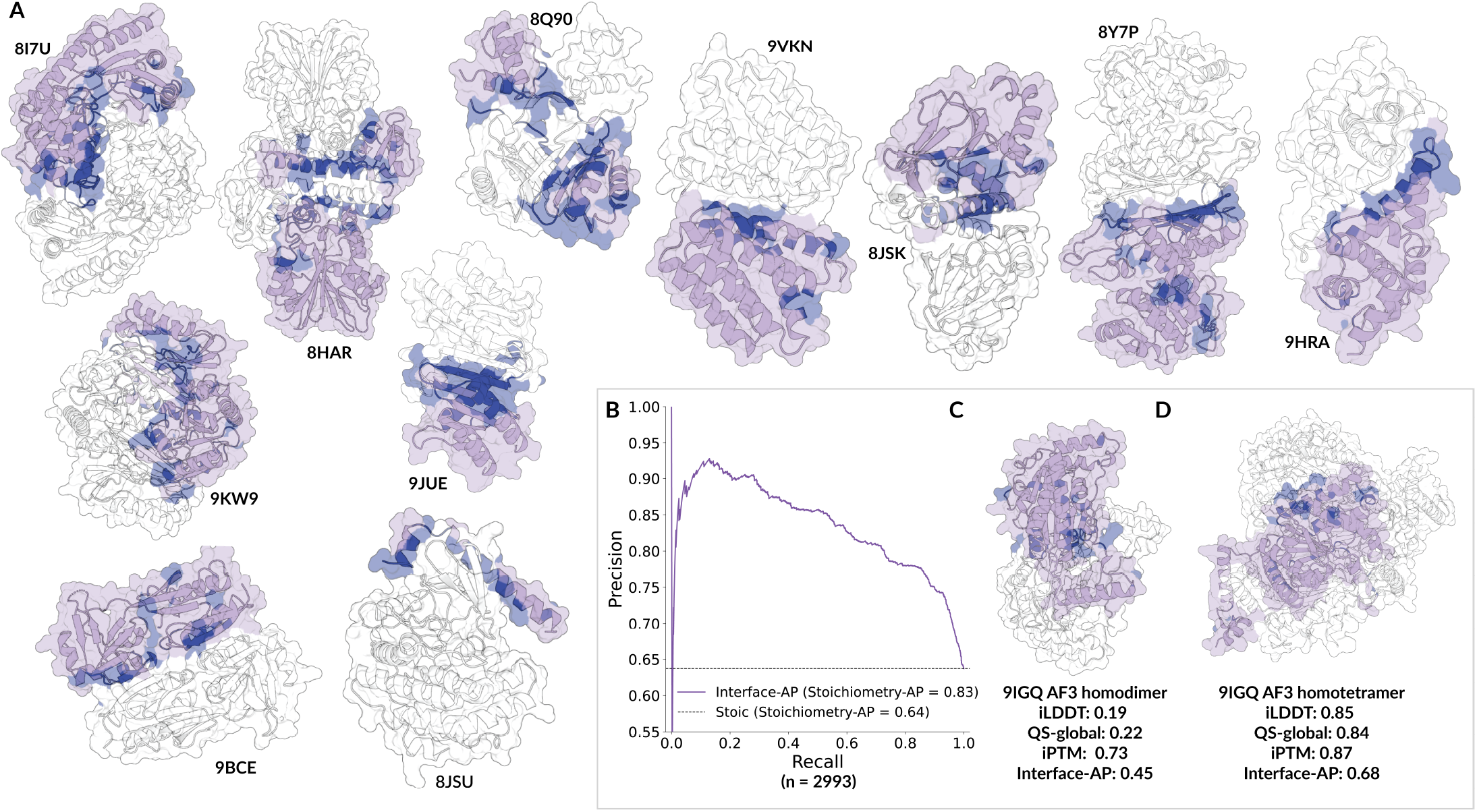
Interface residues relevant for stoichiometry prediction. **A)** Examples of *Stoic* residue weights for one chain each of homo- and heterodimers where all entities have *<*30% sequence identity to the training set. For each dimer, residues with predicted weights *>*0.5 for one chain are highlighted in blue. **B)** Precision-recall curve evaluating Interface-AP as a discriminator between correct and incorrect stoichiometry predictions across the CAMEO dataset. Interface-AP, defined as the average precision between *Stoic*’s predicted residue weights and the interface residues observed in a predicted structural model, achieves an AP of 0.83 for differentiating between correct and incorrect stoichiometries compared to 0.64 (*Stoic*’s accuracy on the non-monomeric set). **C)** AlphaFold3 model of 9IGQ with the top ranked homo-dimer stoichiometry (iLDDT = 0.19, QS-global = 0.22, Interface-AP = 0.45). **D)** AlphaFold3 model of 9IGQ with the third ranked homo-tetramer stoichiometry (iLDDT = 0.85, QS-global = 0.84, Interface-AP = 0.68).

While the *No-Auxiliary* model achieves comparable global stoichiometry accuracy to *Stoic* (Figure 2A), the auxiliary interface loss provides several practical advantages. First, the interface residue predictions offer an interpretable and orthogonal measure of confidence: cases where the model assigns high weights to biologically plausible interface regions are more likely to have correct stoichiometry predictions, enabling users to assess prediction reliability beyond the classification probability alone. Second, the predicted interface residues are themselves a useful output, providing residue-level annotations that can inform downstream applications such as protein-protein interaction or protein design. Third, as shown in Figure 2B, *Stoic* achieves substantially higher precision in identifying interface residues compared to the *No-Auxiliary* model (AP = 0.83 vs. 0.39), confirming that the auxiliary loss is essential for learning biologically meaningful residue weights.

Since *Stoic* shows good performance on the interface residue prediction task from protein sequence alone, we hypothesized that for a correctly predicted stoichiometry, the resulting structural models from AlphaFold3 should place these residues at subunit interfaces, whereas an incorrect one would not. To quantify this, we compute Interface-AP for each AF3-predicted structure between *Stoic*’s predicted weights and the actual interface of the model. For heteromeric targets, we average AP across all unique sequences. We then evaluate whether this score can distinguish correct from incorrect stoichiometry predictions across the non-monomeric CAMEO dataset (for predicted monomers we set AP to 0 since there are no positive residue examples). Using Interface-AP as a discriminator achieves an AP of 0.83, compared to 0.64 (*Stoic*’s accuracy on the subset; Figure 4B), demonstrating that the agreement between sequence-predicted interface residues and the modelled interface is a reliable signal for identifying the correct stoichiometry. Figures 4C and 4D illustrate this for PDB entry 9IGQ: the top ranked homo-dimer prediction of *Stoic* yields a poor model (iLDDT = 0.19, QS-global = 0.22) with a low Interface-AP of 0.45, while the third ranked homo-tetramer produces a substantially higher-quality structure (iLDDT = 0.85, QS-global = 0.84) with an Interface-AP of 0.68. This suggests that Interface-AP can serve as a sequence-derived confidence metric for stoichiometry selection, complementary to structure prediction confidence scores.

## Discussion

Our results demonstrate that *Stoic* successfully addresses the critical bottleneck of stoichiometry prediction in determining protein complex structure. The design of our model makes it easy to integrate with existing structure prediction pipelines - *Stoic* can serve as initial component providing a set of the most likely stoichiometries for multimeric structure prediction methods such as AlphaFold-Multimer (Evans et al., 2021), AlphaFold3 (Abramson et al., 2024) etc. Our evaluation on CAMEO data confirms this in practice: AlphaFold3 models generated with *Stoic*-predicted stoichiometries achieve higher iLDDT and QS-global scores than those built with the naive assumption of one copy per entity, demonstrating that accurate stoichiometry prediction directly translates to better structural models.

There are several promising directions for future development. First, incorporating negative examples during training could further improve model performance. Currently, our training data consist only of positive examples of known protein complexes. Adding negative examples, such as proteins that are known not to interact or form stable complexes, could help the model better distinguish between plausible and implausible stoichiometries. This would be particularly valuable for identifying cases in which proteins might form different complexes under different conditions or where some parts of a complex are unknown. The top-3 global accuracy results suggest that there remains room for improvement in stoichiometry ranking. This could be addressed through more sophisticated ranking algorithms, i.e. by implementing a confidence prediction head.

The interface-aware pooling mechanism developed in *Stoic* has broader implications beyond stoichiometry prediction. Residue-level pooling that focuses on functionally relevant regions could be valuable for many other protein-related tasks, including protein-protein interaction prediction, functional site identification, and drug binding site prediction. The ability to learn which residues are most important for a given task through auxiliary losses represents a generalisable approach that could be applied across diverse problems.

## Acknowledgements

We thank Xavier Robin and other members of the Schwede and Engel groups for technical support and useful suggestions, sciCORE at the University of Basel (https://scicore.unibas.ch/) for providing computational resources and storage space. This work uses outputs from AlphaFold3, subject to the AlphaFold 3 Output Terms of Use (version 3.0.0 [commit 2ffe43f], accessed on 15 Nov. 2024). All AlphaFold3 calculations and analyses were carried out by and using the weights provided to the Schwede group.

---

**Supplementary Figure S1.**
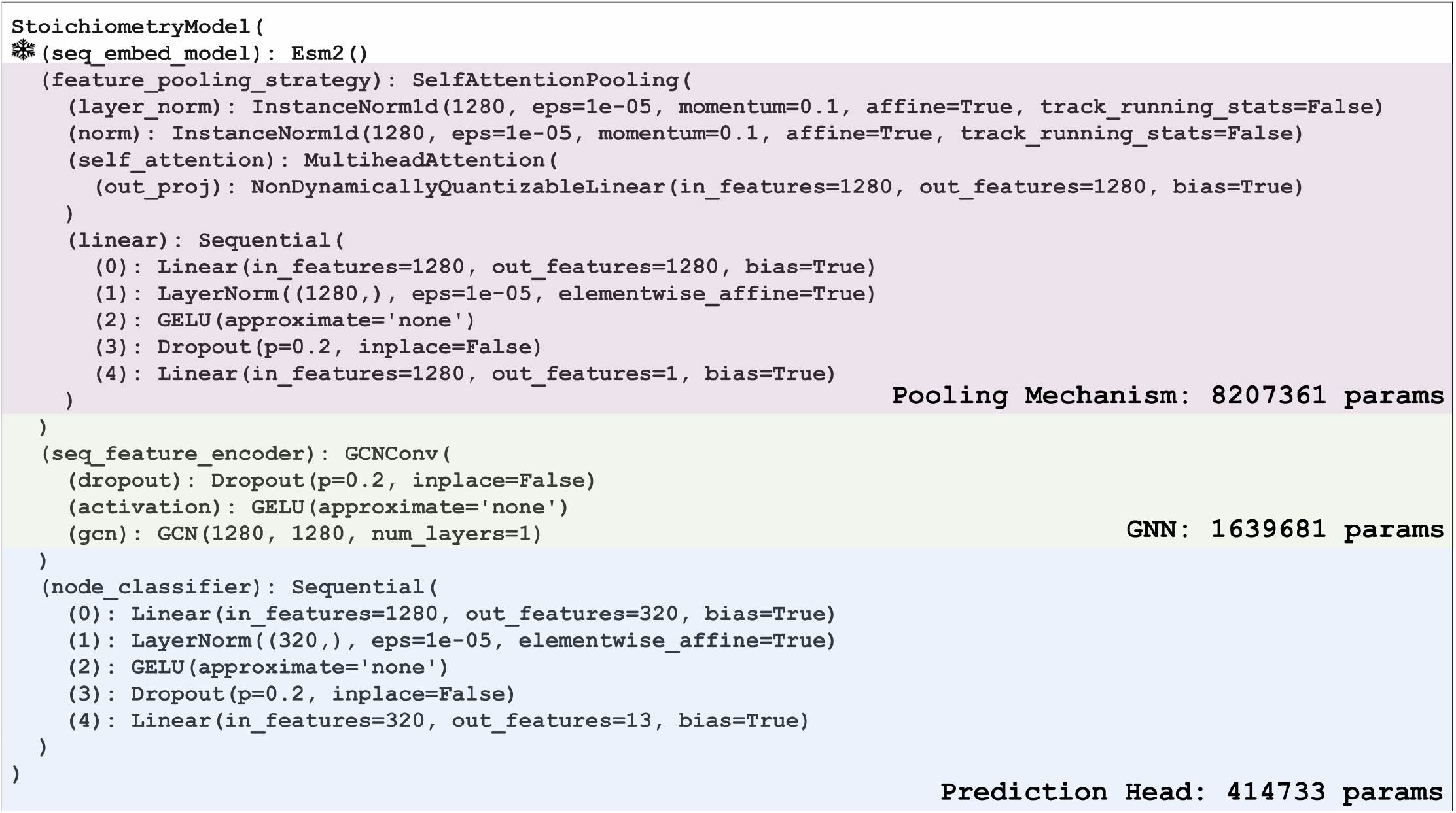
Detailed architecture of Stoic.

**Supplementary Table S1.** Mapping of prediction classes between Seq2Symm and *Stoic*.

| Seq2Symm | Stoic |
| --- | --- |
| C1 | 1 |
| C17 | 17 |
| C2 | 2 |
| C3 | 3 |
| C4 | 4 |
| C5 | 5 |
| C6 | 6 |
| C9 | 9 |
| D12 | 24 |
| D2 | 4 |
| D3 | 6 |
| D4 | 8 |
| D5 | 10 |
| I | 60 |
| O | 24 |
| T | 12,24 |
| CX | undefined |
| DX | undefined |
| H | undefined |
| HOTI | undefined |

**Supplementary Figure S2.**
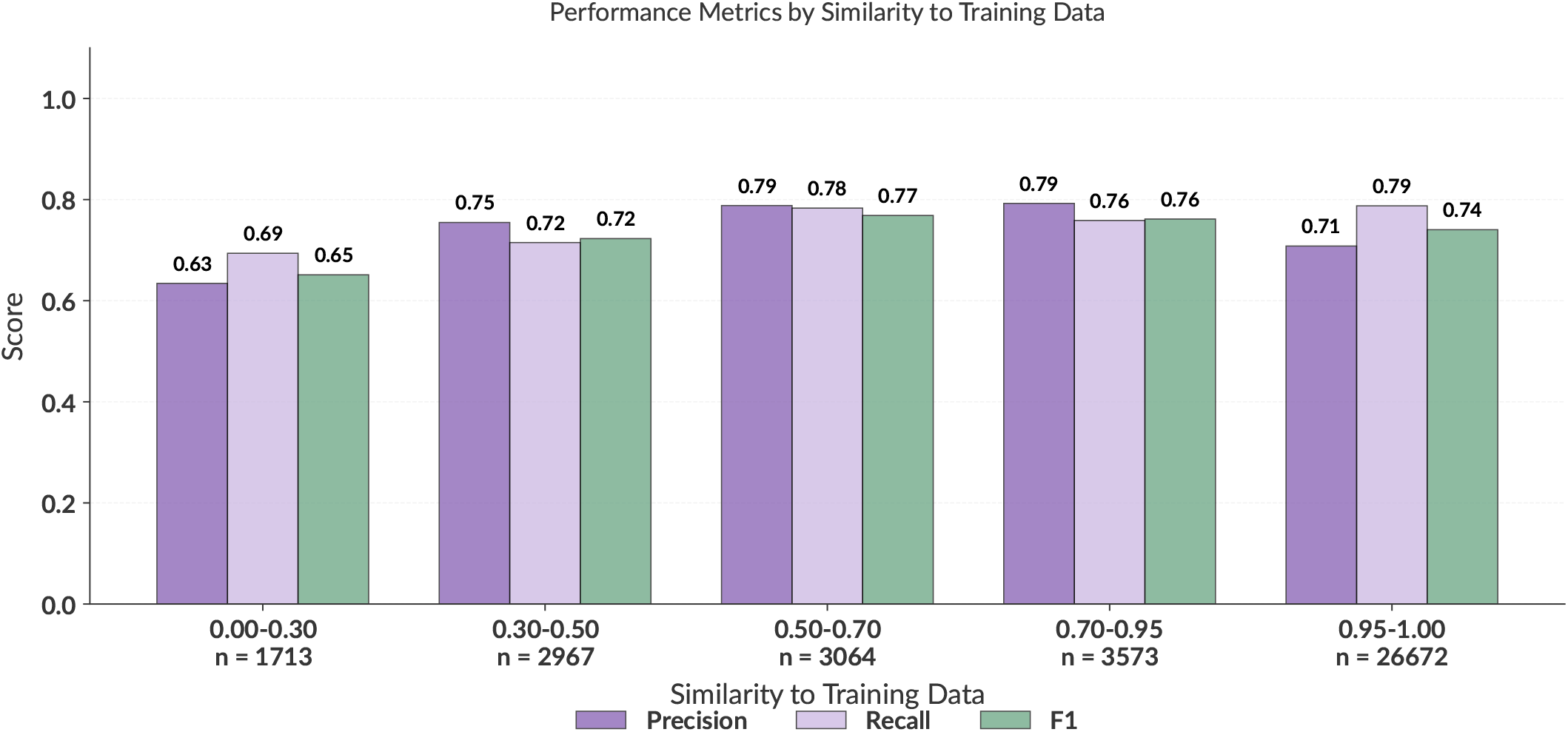
Copy number prediction performance of Stoic depending on similarity to the train dataset.

**Supplementary Figure S3.**
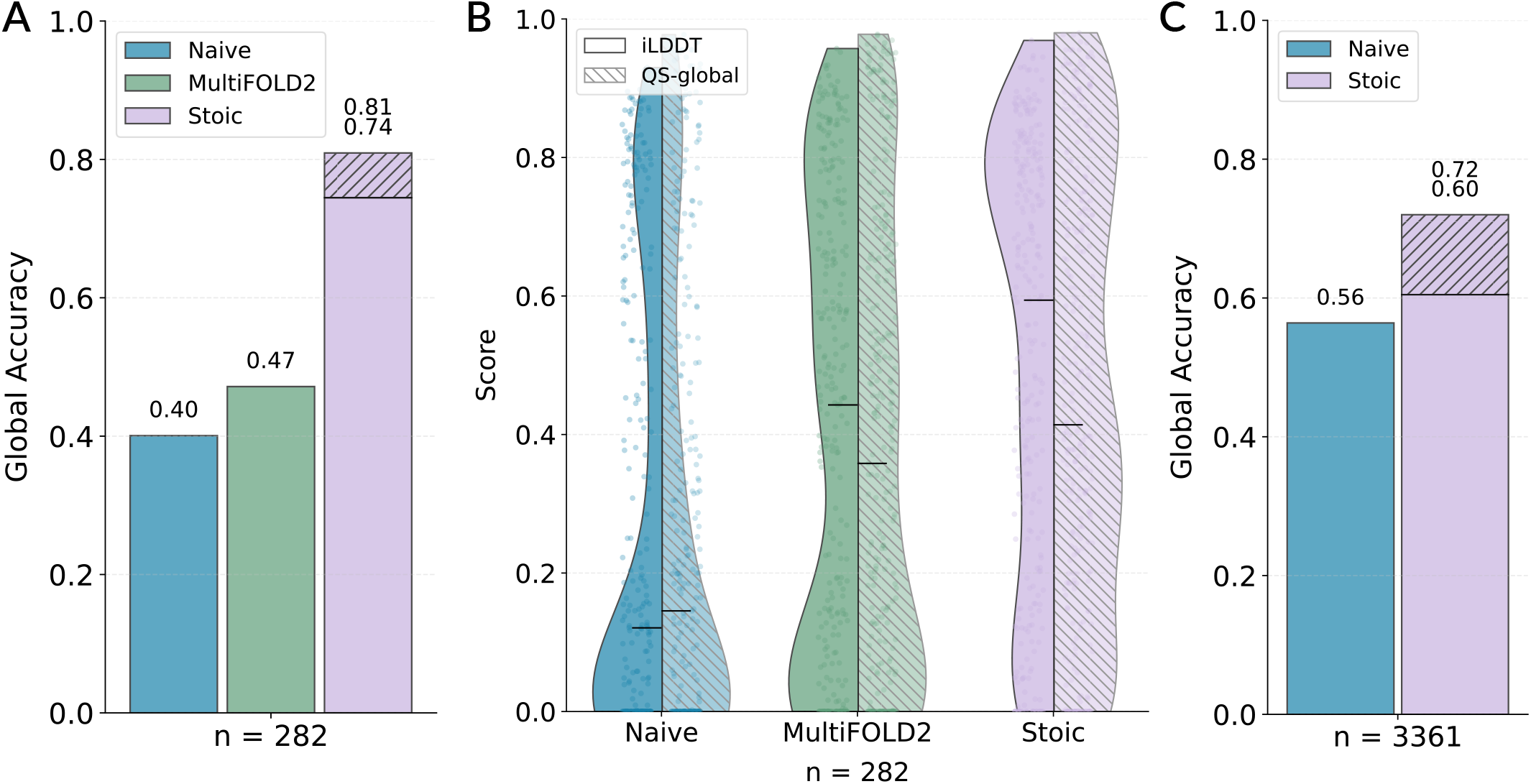
Stoichiometry and structure prediction performance. **A)** Global stoichiometry prediction accuracy on the CAMEO subset for which MultiFOLD2 (green) predictions are available, comparing Naive prediction (copy number of one assigned to every entity) and *Stoic* (purple; best-of-3 shown as a dashed bar). Monomers are excluded because iLDDT and QS-global are not defined for them. **B)** Distribution of iLDDT (solid) and QS-global (dashed) scores for AlphaFold3 models generated using Naive (blue), MultiFOLD2 (green), and *Stoic* (purple) predicted stoichiometries. **C)** Global stoichiometry prediction accuracy on the Complex Portal (Balu et al., 2025) dataset, for which experimental stoichiometry annotations are available.

